# The feasibility of atrial and ventricular arrhythmias recognition using metrics of signal complexity for heartbeat intervals

**DOI:** 10.1101/612002

**Authors:** Ming Huang, Koshiro Kido, Naoaki Ono, Altaf-UI-Amin, Toshiyo Tamura, Shigehiko Kanaya

**Author notes:** Ming Huang is the corresponding author of this paper.

## Abstract

It is well known that the cardiac system is controlled by the complex nonlinear self-regulation and the heartbeat variability (HRV) is an independent indicator of the autonomic regulation. With the assumption that intrinsic differences of atrial fibrillation (A) and ventricular ectopic arrhythmias (V) can be unveiled by a proper approach based on of signal complexity, we examine the feasibility of detecting these arrhythmias of different pathological origins using metrics of complexity for heartbeat intervals (HRI). Specifically, the normal sinus rhythm (N), the A type and the V type are used as the targeted types of heartbeat. By extracting the entropy-based features from HRI of different lengths, i.e., from 300 heartbeats to 1000 heartbeats, we examined the distinguishability of these 3 types of heartbeat. By applying the features to the random forest model, the HRI signal of 600-heartbeat-length can be used to detect the A and V completely, i.e., with 100% of type-wise recall and precision. What is more, this approach is sensitive to the existence of the corresponding arrhythmias. The results substantiate our assumption about the intrinsic difference of the A and V type. A further investigation applying this approach to a wider spectrum and a finer stratification of arrhythmias/ cardiac diseases and may lead to the systematic understanding in the context of complexity and better insight for its practical use for wearable/unconstrained monitors.

## 1. Introduction

It is well known that the cardiac system is controlled by the complex self-regulation (Narayanan *et al* 1998), involving various physiological indices such as blood pressure, body temperature etc. Moreover, the nonlinearity of the regulation may be attributed to the complexity of the fractal-like structure of the His-Purkinje fiber network of the heart (West *et al*, 1999). It is plausible that defects of the afferent system or electrical transduction system may give birth to the difference in terms of complexity. On the other hand, the cellular/molecular defect of the cardiomyocyte may also be manifested as the cardiac arrhythmias (Hong *et al*, 2014) and expressed by the change of the complexity.

Measures that were defined based on entropy are appropriate to measure the complexity of signal and a series of modifications and validations for entropy-based metrics have been made. Among these measures, sample entropy (SampEn), which exclude the self-match, is a modification over the approximate entropy and have been used to analyze the heart rate variability (Richman and Moorman, 2000). The SampEn requires 10^*m*^∼30^*m*^ (*m*: the length of a compared run of data) and lacks consistencies in some cases. To amend the issues of SampEn, Costa *et al* (2002) proposed the multiscale entropy (MsEn), in which the original signal is coarse-grained by moving average over different scales without overlapping and the entropy values are calculated for each scale. It enables a deeper investigation of the signal of interest with multi-resolution and gives more stable results (Costa *et al* 2005). However, it requires a relatively long signal. More recently, the refined composite multiscale entropy (RCMsEn) has been proposed to address the issue of signal length by Wu *et al*. (2014).

It has become more clear that many cardiac arrhythmias can be characterized on the basis of the physical principles of nonlinear dynamics (Christini *et al* 2000). Researchers have been trying to characterize arrhythmias of different origins so as to distinguish them. Owis *et al* (2002) succeeded in distinguishing the normal sinus heart rhythm from the abnormal rhythm using the correlation dimension and the Lyapunov exponents. Zhou *et al* (2014) have tried to use the Shannon entropy in detecting the atrial fibrillation from normal sinus heart rhythm and have shown promising results in a real-time application.

The measurement of the heartbeat interval (HRI), which is canonically defined by the R-R interval of electrocardiograph (ECG), is appropriate to reflect the cardiac self-regulation. Moreover, HRI can also be obtained by other techniques, e.g., Plethysmograph or even the Ballistocardiogram. The diversification of the signal source endows it a much broader field for applications, from the clinical setting to personal healthcare setting (Krivoshei *et al* 2017). The progress in wearable devices and IoT in recent years give this approach that analyzes the complexity of cardiac system self-regulation based on HRI (abbreviated as ACCaHRI hereafter) a much more clear prospect.

The wearable/unconstrained modalities provide more flexibility and more dynamic information at the expense of lower signal quality due to the improper measurement setting or body movement etc (Huang *et al* 2017, Tang *et al* 2017). Sometimes, the signal is so severely contaminated by the noise that only the heart rate information (e.g., from the R-R interval) can be extracted from the ECG signal. Hence, a reliable approach using the metrics derived from HRI alone to reflect the physiological/pathological status of the heart is a practical necessity (Chen *et al* 2011, Jarchi *et al* 2017).

Recent studies (Zhou *et al* 2014, Krivoshei *et al* 2017) have shown a looming picture of distinguishing the cardiac problems/arrhythmias based on ACCaHRI. However, from a holistic point of view, a general picture about the projection of the cardiac problems/arrhythmias of different origins onto the entropy features space is indispensable to evaluate this approach and make necessary amendment.

Specifically, a binary classification, e.g., detection of abnormal from normal, would be grateful if one wants to simply know one’s physiological condition. However, for a more detailed pathological diagnosis, a multi-classes classification in the feature space is necessary.

After the summarization of the achievements and issues of the ACCaHRI, we propose our assumption and aim of this study as follow:

### Assumption

For the arrhythmias originate from different chambers of the heart, the autonomic regularization takes different measure to compensate for their influence. A proper measure of signal complexity is adequate in projecting the HRIs of normal sinus rhythm (N) and arrhythmias of different origins, i.e., the atrial fibrillation (A) and ectopic ventricular heartbeat (V) onto a new feature space so that they become differentiable in that space. We choose the A and V because that these 2 arrhythmias originate from different physiological tissue, i.e., the atria and the ventricles, and that they are relatively common, which ensure the necessary amount of the HRI samples for analysis and the training of machine learning model.

### Aim

By using a proper measure of complexity in combination with machine learning algorithms, we can distinguish the N, A, V with accurately and precisely with a relative short HRI signal (80 bpm, about 6 ∼ 12 mins). The reason to emphasize the signal length lies in that a short signal would be easier to obtain by a wearable device.

This study contributes to the initiative holistic exploration of the possibility of using the ACCaHRI to provide detailed physiological/pathological information targeting especially at the wearable devices. Attempts applying a wider spectrum and finer stratification of arrhythmias could be anticipated.

## 2. Materials and Methods

The approach in this study was based on the statistical measure of the HRI, therefore, it tried to detect the targeted arrhythmias in a short time interval. The A and V were used as the targeted arrhythmias in this study because there are relatively common and physiologically significant on the one hand. On the other hand, they are distinguishable in a feature space based on our assumption.

In this study, the recently developed RCMsEn which is an improved measure based on multiscale entropy was integrated with a nonlinear ensemble machine learning method, the random forest, to project the HRI time series onto a nonlinear feature space and to divide the space for heartbeat classification. This integrated method aimed at facilitating the use of wearable measurement of HRI as a diagnostic assistant.

### 2.1 Data and preprocessing

We chose the MIT-BIH arrhythmia database (MITDB) (Goldberger *et al* 2000) with manual annotations of the heartbeats from the well-established Physionet, which is a data hub for physiological signals. In the database, there are 48 records of 360Hz-sampled ECG signals including a variety of arrhythmic ECG signals. Since each record in this database has been annotated by two physiologists for each heartbeat in view of the ECG morphology of the waveform, we use the annotations as the reference of the signal. Moreover, the annotations have been adjusted to the R-wave peaks, and they are accurate enough as the reference in HRV studies (Moody and Mark, 2001). Hence, the HRI were extracted based on the R-peak annotations in this study.

The correspondences between the 3 targeted heartbeat types used in this study and the rhythms defined in the MITDB are:

N: N; A: AFIB; V: B and T.

The annotations after each colon are the counterparts defined in MITDB, whose full names are sinus rhythm, atrial fibrillation, ventricular bigeminy, and ventricular trigeminy. Single ventricular premature beat was not included due to its relatively low density in a short signal segment. With regard to the labeling of a segment, heartbeat density (*σ*) criteria as follow were used:

N: *σ*_N_ ≥ 0.95; A: *σ*_A_ ≥ 0.7; V: *σ*_V_ ≥ 0.15.

The justification of this setting will be discussed in the Discussion section. We labeled the segment as A or V if and only if the corresponding heart rhythm satisfies the criteria above; The HRI was extracted as the time intervals between the R-peak of 2 consecutive heartbeats.

### 2.2 SampEn, MsEn and RCMsEn

The RCMsEn is considered as an appropriate measure to reflect the complexity of an HRI signal because it provides multi-resolution information and works well for a relatively short signal. The relevant SampEn and MsEn will be introduced briefly to facilitate a quick grasp of RCMsEn.

Suppose there is a short signal *X* = (*x*_1_, *x*_2_, …, *x*_*n*_), the SampEn is defined by

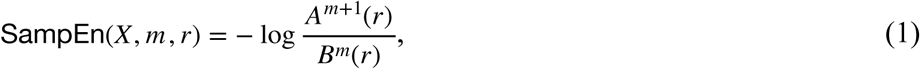

where *m* is the embedding dimension, which is set to 2 in most of the cases; the *r* is the tolerance and it is usually set as 0.1∼0.2 of the standard deviation of the signal. According to the setting of *m*, the signal is divided into fragments of a length of *m* and *m*+1. The *A*^*m*+1^(*r*) is the number of pairs 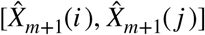, (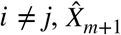: the fragment of *m*+1 length), whose distance is shorter than *r.*

The MsEn was proposed to provide multi-resolution information based on the entropy theory, in which the original signal is averaged over different scales defined by τ (scale *k* = [1,*τ*]). Therefore, for each scale *k*, a new time series is generated as

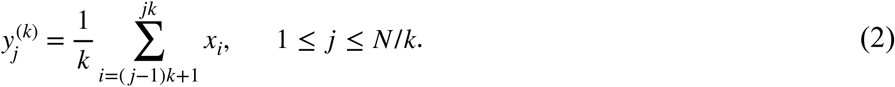

For each scale, the generated time series *y*^(*k*)^ is used to calculate the SampEn. Hence,

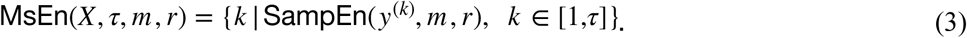

The entropy values of each scale together give us the chance to inspect the self-similarity of the signal from different time scales. However, the coarse-graining of the MsEn would decrease the length of the signal generated by a ratio of the scale.

RCMsEn is a modification based on the MsEn to solve the issue of data-length for the short signal, which generates *k* new time series 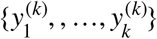 for the corresponding scale *k* by sifting the coarse-graining procedure to the right as shown in Figure 1 (b).

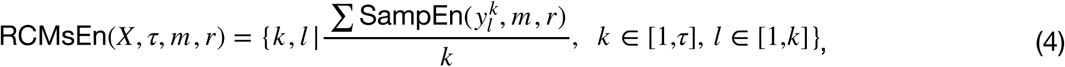

where the SampEn values are calculated individually and averaged over *k* for each scale. It has been validated that the RCMsEn would provide more stable and consistent values, for especially short time series (Wu *et al* 2014).

**Figure 1.**
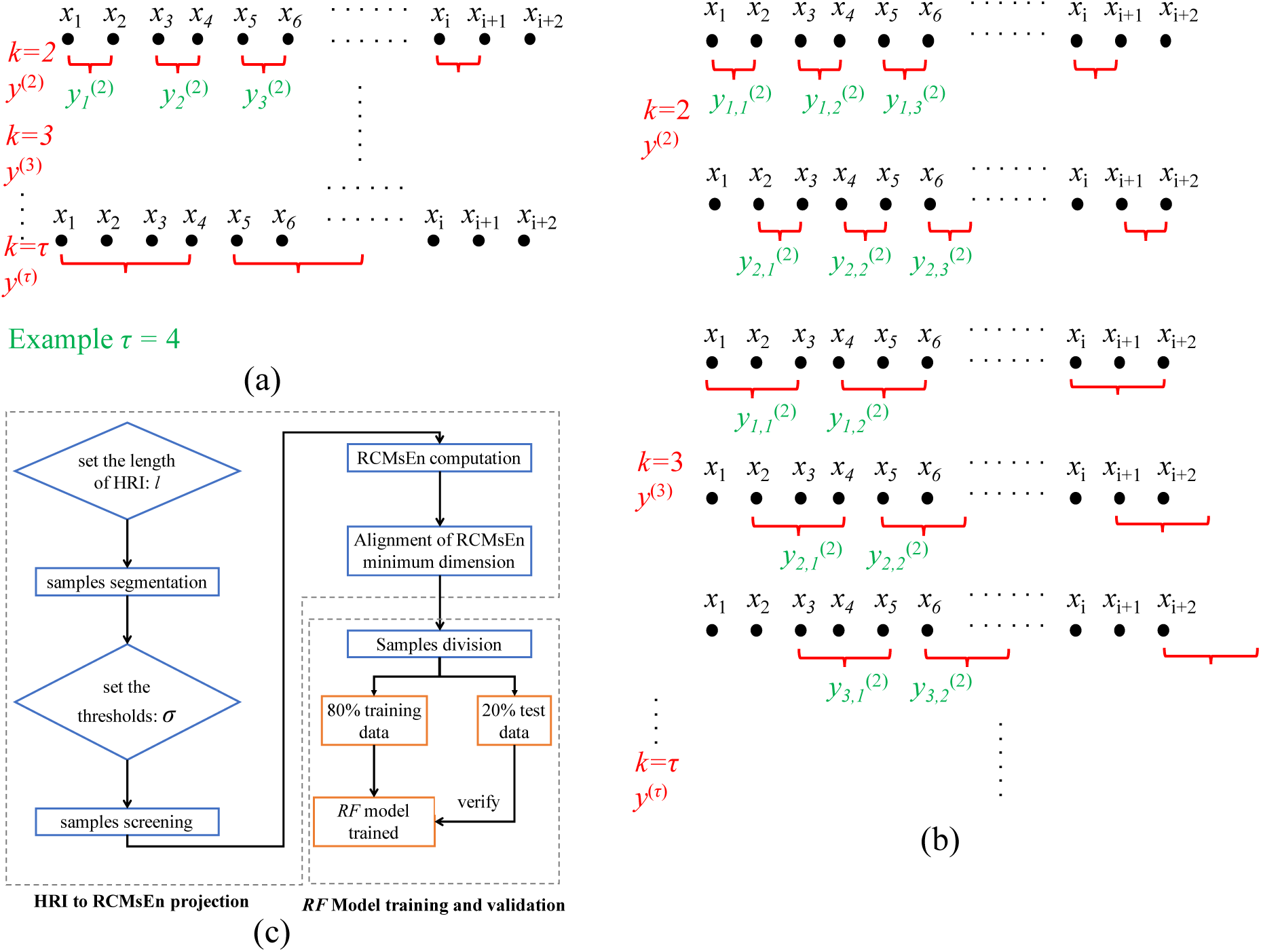
Illustration of the MsEn (a) and RCMsEn (b). MsEn provides multi-resolutions information by averaging the raw signal over different scale *k, k* = [1,*τ*]. (c) is the flow of the approach.

### 2.3 Comparison of different data-length

As it is introduced above, the length of the HRI signal (termed as data-length hereafter) is of interest in this study, we generated the RCMsEn values for HRI signal of different data-lengths, whose range were [300,1000] with 100 intervals. The corresponding time lengths were about 4∼12 minutes (80 bpm).

The scale parameter τ was set as 20, which means that for each segment of different data-length, the maximum dimension of the output from the RCMsEn is 20. Noteworthily, the dimension for the short signal may decrease due to the undefined entropy, for this, the dimension of the output for all the 3 types of heartbeats, i.e., N, A, and V were aligned to the minimum dimensions of these 3 types.

Noteworthily, even for the same type of heartbeat, the dimension may vary over samples, the maximal scale was chosen in a way that if more than 1% of the samples showed undefined entropy, the scale was excluded.

### 2.4 The classifier: Random forest

The random forest (*RF*) is an ensemble learning algorithm, which by growing a number of tree-based weak classifiers to prevent the overfitting problem that can often be seen in a single complicated model. At the same time, the *RF* reduces the predictive variance by decorrelating the individual weak classifier by the random selection of partial independent variables to grow a tree, and by growing it with different bootstrapped samples.

The RF was chosen based on the analysis of the RCMsEn values. We applied the principle component analysis to the RCMsEn values (results not shown in this paper), whose result showed that the 3 types of heartbeats cannot be separated linearly. The difficulty in applying linear methods may be due to the high variances in some features. *RF* was chosen because it is adequate in decreasing the variance in the prediction by averaging over the individual trees. The RF was implemented as follow:

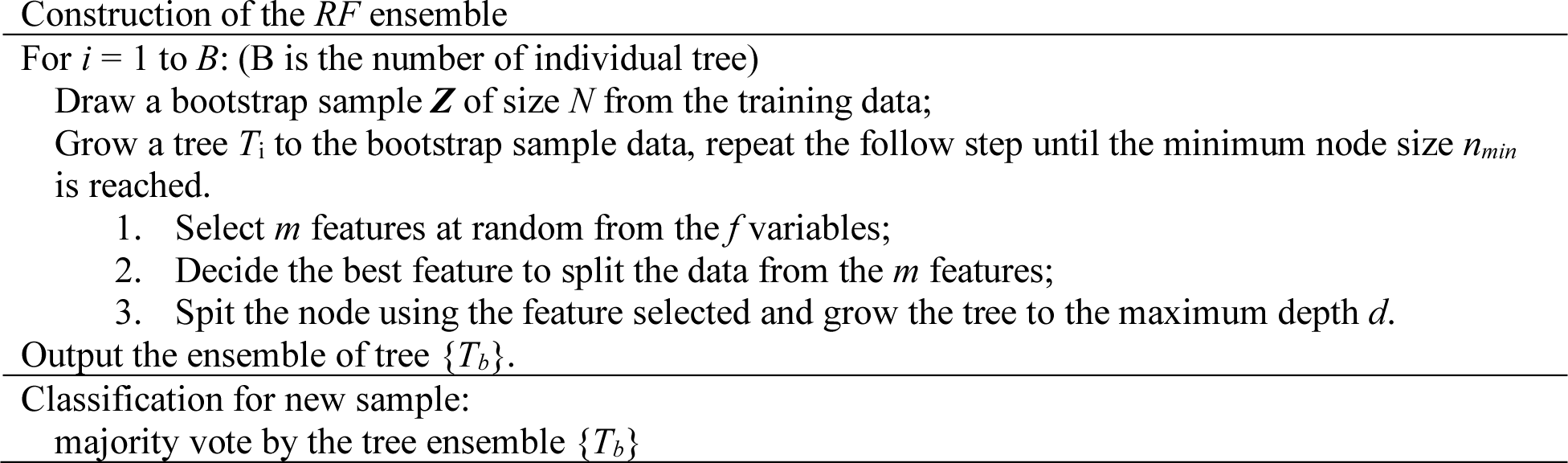

The *RF* reduces the variance of the of the ensemble according to the following equation

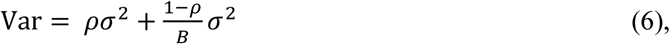

where the ***ρ*** is the correlation of trees and ***σ***^**2**^ is the variance of the features (assumed as a constant for all features). By increasing the tree numbers the second term on the righthand side becomes minor; while the first term can be decreased by reducing the correlation of trees by a random selection of the *m* features.

The *RF* also shows its advantage in the model interpretation. Unlike the popular deep learning approach, where the importance of each independent variable is difficult to keep track of, the importance of a variable can be reflected by averaging its importance over each tree grown. The variable importance is evaluated by averaging the decrease in accuracy over all the individual trees when we permute the variable of interest.

The hyperparameters for the *RF* are the *number of trees, the max features* and the *max depth* of a tree. If the computation time is not the parameter of concern, the larger is the tree number (100 in this study) the smaller is the variance. As is mentioned above, only a portion of the independent variables (features) are used to grow a tree, and the number is defined by the *max features* 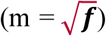. Finally, a tree with a deep depth may cause the overfitting problem, therefore, it is necessary to define the *max depth* of a tree (*d* = 10 in this study).

### 2.5 The flow of the approach and Evaluation of the Results

The flow of this study from the HRI extraction to *RF* model construction and model validation is shown by Figure 1(c). The global hyperparameters are the length of the HRI and the thresholds of density for the three heartbeat types in one segment.

The workflow introduced above will conceptually split the feature space consisting of the RCMsEn values for the 3 targeted types of heartbeat, which can be instantiated by a machine learning model. To train and test the model, training samples and test samples are split at a ratio of 4:1 for each type of heartbeat. To evaluate the combined performance of the RCMsEn-based features and the *RF*-based classifier, class-specific precision and recall are used.

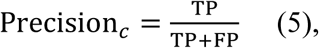

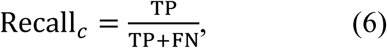

where subscript *c* denotes the class-wise calculation.

A deeper understanding of the features of different scales on the classification can be revealed by the analysis of the importance of the *RF* model. For each data-length, the importance of each scale can be obtained from the *RF* model. For the first 9 important scales in terms of median, the *Mann-Whitney U test* is used to test the significant difference between 2 scales in view of the small number of samples.

## 3. Results

### 3.1 Features based on RCMsEn

The situation that a sample of relatively short data-length may cause the undefined entropy still exist. The scales available for HRI of different lengths are tabulated in Table 1.

**Table 1.**
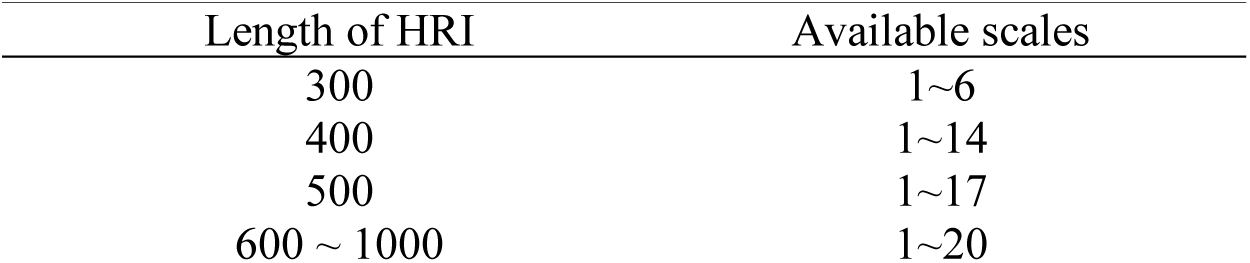
Available scales of RCMsEn for HRI of different lengths

| Table 1 Available scales of RCMsEn for HRI of different lengths |  |
| --- | --- |
| Length of HRI | Available scales |
| 300 | 1~6 |
| 400 | 1~14 |
| 500 | 1~17 |
| 600 ~ 1000 | 1~20 |

The overall trends of the 3 kinds of heartbeats are displayed by the boxplots in figure 2, in which samples of 600 data-length are used. From the figure, it can be seen that the decreasing trends along the scales are similar; whereas indistinct differences of the numerical distributions are profiled.

**Figure 2.**
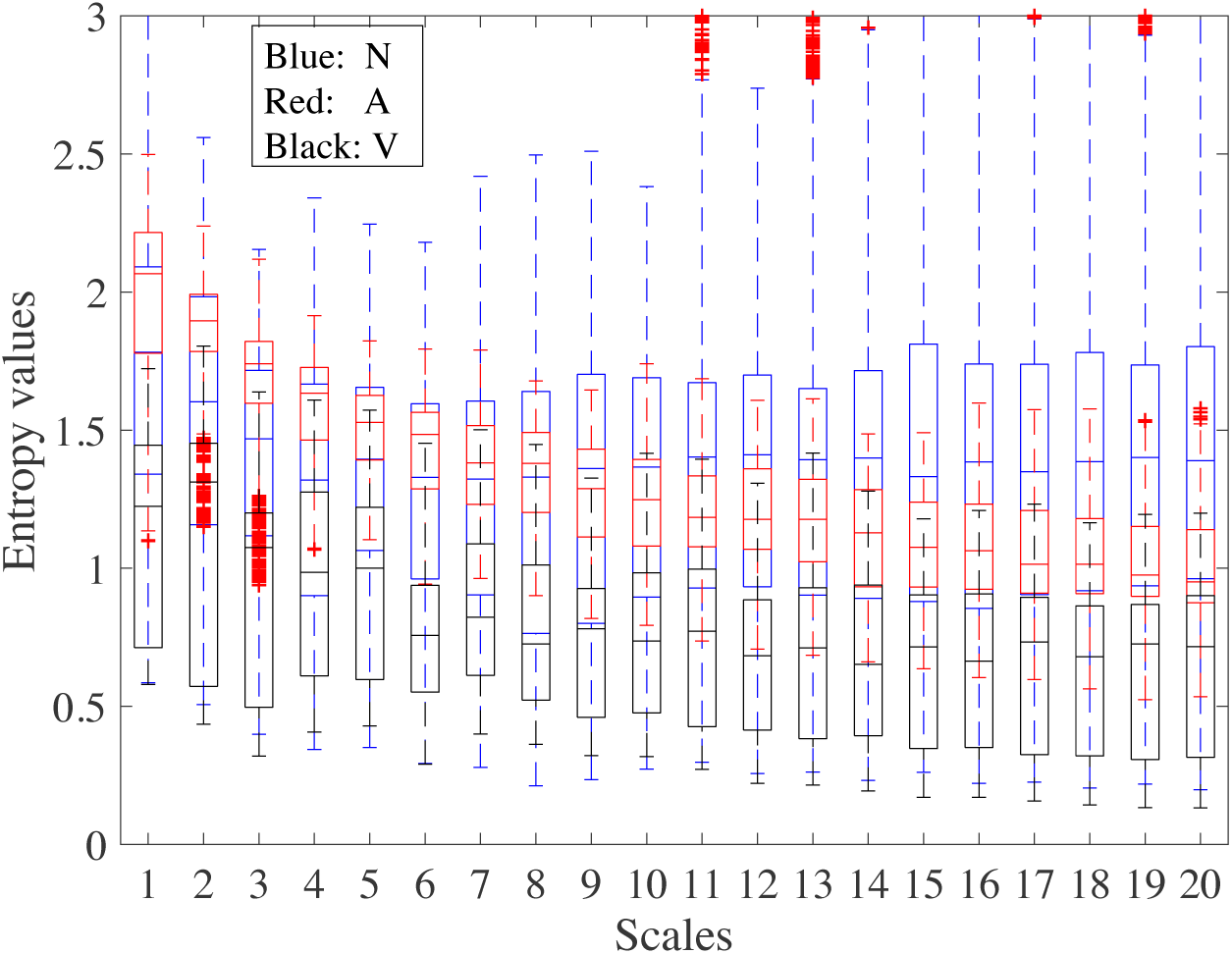
Box plots of the RCMsEn values of segments with 600 data-length over the 20 scales. Blue boxes represent the N type; Red boxes the A type and Black boxes the V type.

The A type has the largest median values for the first 7 scales and V type has the smallest median values over all scales. It can also be seen that the variance turns bigger along with the increase in scales, the situation is especially phenomenal when the scale is larger than 10.

### 3.2 Recall and precision of the RF model

The dimension of the RCMsEn features increases with the data-length as shown in Table 1. The effect of the increase in the dimension of features can be reflected by the performance of the *RF* model. In Figure 3, the recalls and the precisions for all 3 types of heartbeat increase significantly from 300 to 400 data-length, among these improvements, the situation is more so for the A type. The *RF* model reaches the status of perfect detection when the data length comes to 600. It has a better performance when compared to the study of Jovic and Jovic (2017). By combining some HRV additional features with the alphabet entropy, their results show 83.7% recall and 96.7% specificity for A type; 65.2% averaged recall and 99.3% averaged specificity for V type.

**Figure 3.**
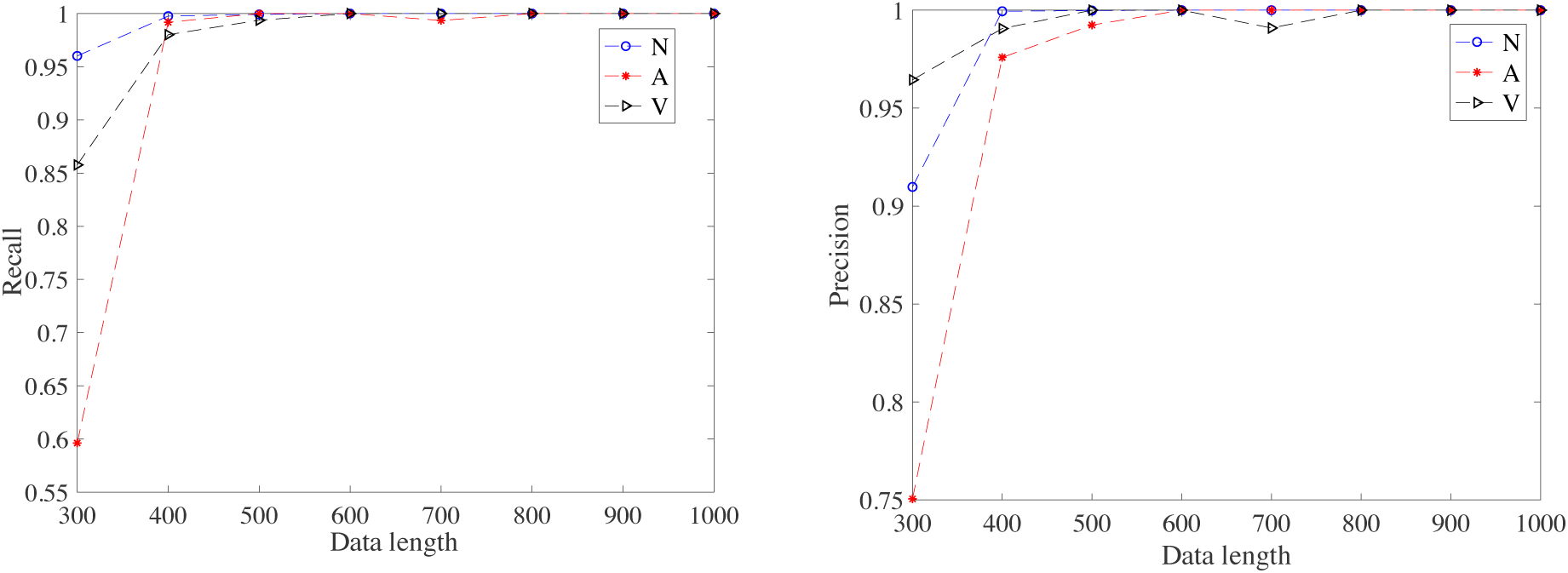
Recall and precision of the RF model using the RCMsEn values as the features. Both the recall and the precision values increase sharply when the length of the HRI increases from 300 to 400.

For the situation when the data length is 300, 37.4% heartbeats of A type and 11% heartbeats of V type are mistakenly classified as N, whereas 3% of N type heartbeats are classified as A type. Moreover, the misclassification between the A and V are not as significant, 3% of V type heartbeats are classified as A type; whereas 3% of A type heartbeats are classified as V type. These results suggest that the N type cover a larger portion of the feature space than A and V when the dimension of the features is insufficient; whereas the intrinsic difference of A and V can be reflected in a feature space of lower dimension.

### 3.3 Importance of the features

The medians of normalized importance of the features are plotted In figure 4, from which it can be seen that the importance of a lower scales (τ < 8) are generally higher and the scale 4, 1 and 3 are the 3 most important scales for the classification.

**Figure 4.**
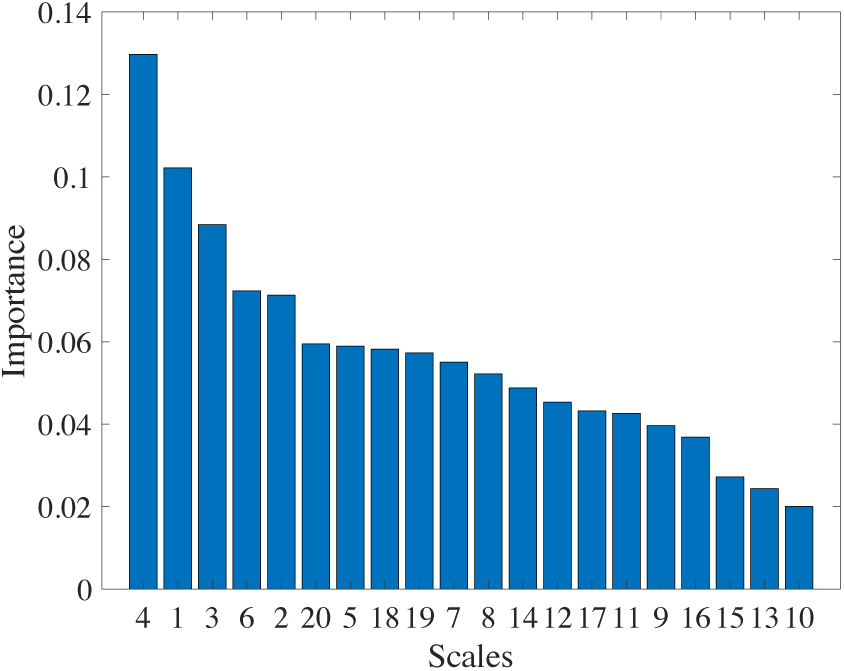
The medians of the importance of features over 20 scales. The x-axis is the scale τ; whereas the y-axis is the medians of the normalized importance.

*Mann-Whitney test* was used to test the differences between the importance of 2 scales for the first 9 important scales, whose accumulated importance is 0.63. The results of tests show that:

- the most important scale 4 is significantly different (*p* < 0.05) from all other scales;
- second important scale 1 is significantly different (*p* < 0.05) from the other scales except for the sale 3, which is significantly different (*p* < 0.05) from the scales that are less important than itself.

## 4. Discussion

The HRI-based heartbeat classification stands out in the applications for wearable and unconstrained measurements of the cardiac electrical signal, whose quality is inferior and prone to be noisy. However, a signal entropy value in the original scale is not adequate in separating different heartbeats. As a follow-up experiment based on the algorithm of Zhou *et al* (2014), we have used the algorithm in this multi-classification problem. From the experiment we got a similar result for the binary classification of N and A by setting the threshold of entropy value 0.63; however, the single entropy value becomes insufficient when the V type was added in. Figure 5 shows the distribution of the Shannon entropy value of each type of heartbeat, from which it can be seen that the V type sprawls along the x-axis and it makes the classification improbable.

**Figure 5.**
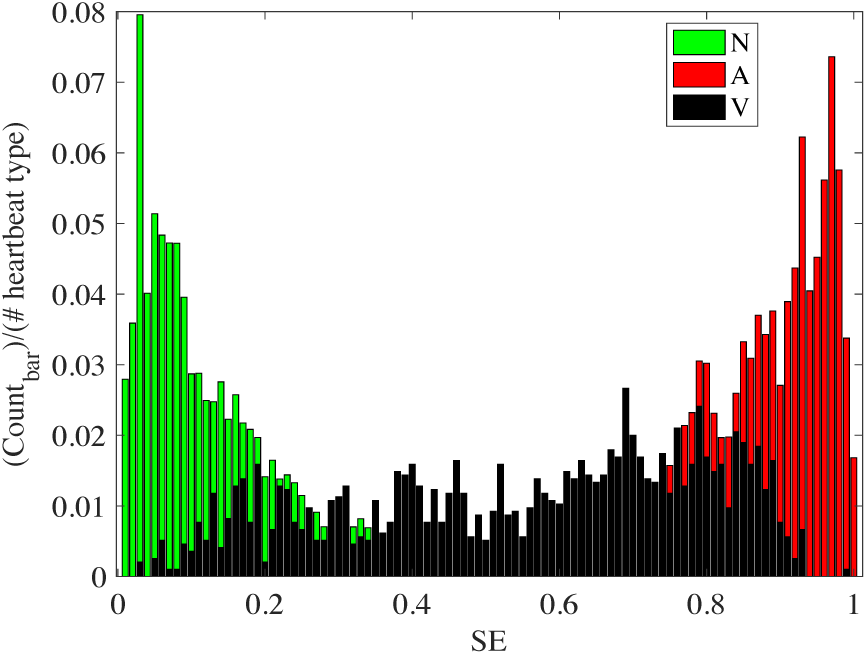
The entropy value of N A and V using the single entropy value. The x-axis is the Shannon entropy value and the y-axis is the proportion of the count number of each bar over the total number of each kind of heartbeat.

There are some hyperparameters to set in our study. A parameter that should be mentioned is the threshold for heartbeat density. The determination of σ_A_ and σ_V_ are based on the preprocessing of the A and V types trying to preserve the samples as much as possible. Therefore, by decreasing the of σ_A_ and σ_V_, the sample number will increase, and the concern now becomes will the decreases density cause a lower recall and precision? We try the grid comparison by setting σ_V_ in [0.10:0.70] with 0.10 interval; σ_A_ in [0.05:0.20] with 0.05 interval, which will generate 28 combinations of density thresholds. Noteworthily, greater portions of A and V samples have a higher density than the ranges we set above. For example, for the 500 data-length, there are 15% of A samples in the range of [0.10:0.70], while other A samples have a higher density than 0.70. Similarly; there are 40% of V samples in the range of [0.05:0.20], while other V samples have a higher density than 0.20. The 300 and 500 data-length are used for the comparison, since their results may be more sensitive to the density according to the classification results. The recalls and precisions of 300 data-length plotted by surface plots are shown in Figure 6, based on which neither recall nor precision shows propensity with the threshold values. Furthermore, the situation for 500 data-length is similar to that of the 300 data-length, but with higher recall and precision values (above 0.95). Since the results for A and V types are close to each other which will cause unclear images, they are omitted here. These comparisons suggest that this approach is sensitive to the existence of corresponding arrhythmias.

**Figure 6.**
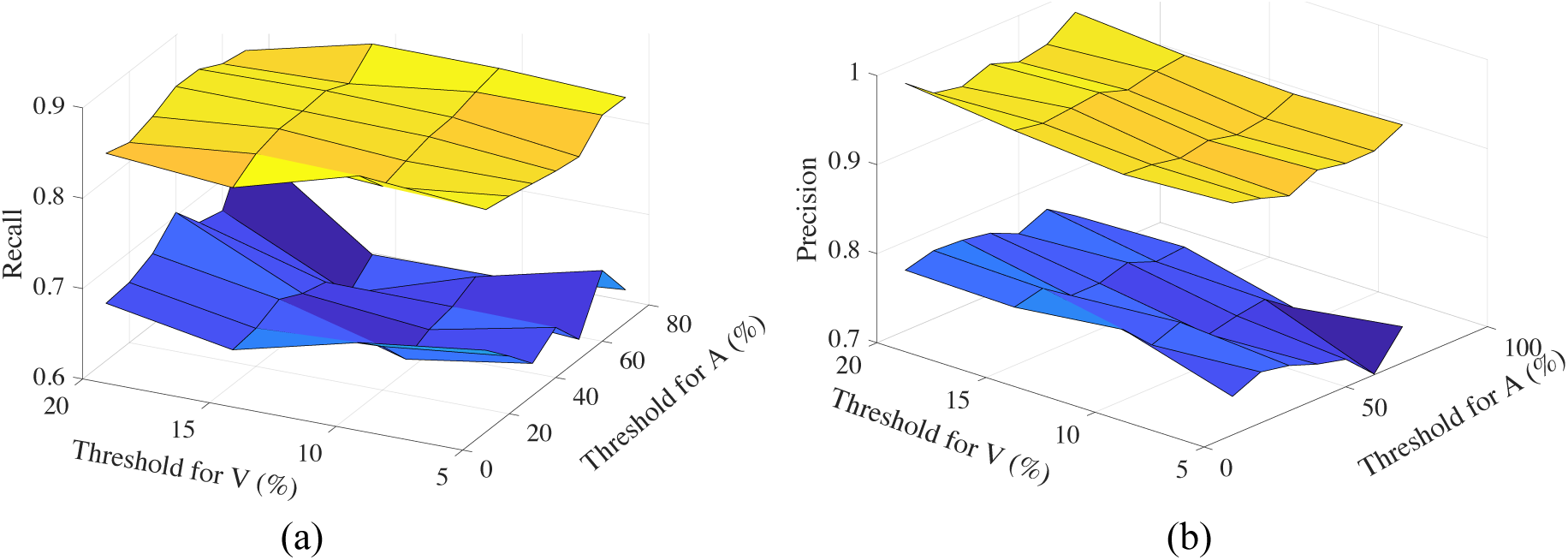
The recalls (a) and precisions (b) of different combinations of density thresholds for A and V type classification for the sample of 300 data-length. Blue surfaces show the results for A type; whereas yellow surfaces show the results for V type.

The results of the RCMsEn computation and the multi-classification based on *RF* model substantiate the underlying assumption about the intrinsic difference of A and V. Similarly, the assumption can be extended to other types of arrhythmias with different pathological origins. In this study we only analyze the samples from the MITDB in considering the amount of annotated data, the following study with a wider spectrum of arrhythmias/cardiac problems would be very interesting.

As we have mentioned in the [Materials and Method] section that the A and V are relatively common. Therefore, we have enough data to train the machine learning model. It is plausible that this method can be extended to a broader range with more arrhythmias with sufficient sample data.

The RCMsEn is an improved version of the multiscale entropy that can generate the entropy values for a relatively short signal. As we can see from the overall profiles of the 3 heartbeat types in Figure 2, some RCMsEn values are of high variances although the clear trend for each type can be seen. In view of this characteristic, by integrating with the nonlinear supervised learning machine *RF*, the RCMsEn can be used in the multi-class classification. More importantly, this integrated method proposes the idea to project and use the HRI in a nonlinear feature space of high dimensions, which additional heartbeat types can be projected to and a systematic study can be performed on. However, as a statistical method, the RCMsEn needs the HRI of a certain length (600 heartbeats in this study), which makes it inappropriate to the detection of some malignant symptoms that need prompt treatment, such as Torsade de pointes.

Based on the results in this study, it is probable that the proposed method is adequate in providing a coarse identification (on the chamber level) for the origin of the arrhythmia. Although AF and ventricular ectopic beats pose no immediate threat to health, long-term evidence is useful in tracking the heart condition, since the AF causes intracardiac blood stasis which may trigger a stroke (Page *et al*, 2003). Ventricular ectopic beats are thought to be relatively benign in the absence of structural heart disease, however, its frequency is associated with mortality in ischemic heart disease. It is also suggested that with a high PVC prevalence, one should pay attention to the progression of the left ventricular dysfunction (Niwano *et al*, 2009).

A hierarchical stratification using this method could be carried out with more reliable data. Medical record digitalization and opening are advocated here for a better study on a systematic level. By taking more arrhythmias into account, the nonlinear nature of the cardiac system which can be manifested by entropy-based measures would become more clear. A number of HRV measures have been proven to be useful in describing the cardiac autonomic regularization (Behar *et al*, 2018), with the support of sufficient ECG records, a systematic study about the manifestation of cardiac automatic regularization based on HRV measures is significant and indispensable as future work.

As we have mentioned, this method is targeted at the HRI extracted from wearable/unconstrained sensors, whose signal may be contaminated by noise. Therefore, the analysis of higher order spectra which is a high signal-to-noise ratio domain may be useful in profiling the difference between heartbeat rhythm nonlinearly.

## 5. Conclusions

In this study, with the assumption that intrinsic differences of atrial fibrillation and ventricular ectopic arrhythmia can be unveiled by a proper method based on signal complexity and nonlinear machine learning model, we propose the approach combining an improved multiscale entropy with the nonlinear random forest model to distinguish the arrhythmias from the normal sinus rhythm using a relatively short data. Our approach shows that with data of 600 consecutive heartbeats, the arrhythmias and the normal rhythm can be distinguished completely. Furthermore, this approach is very sensitive to the existence of arrhythmias. By using this approach, a further study with a wider spectrum of arrhythmias of different pathological origins may be of great values for a systematic understanding of the arrhythmias based on complexity and facilitate the use of heart rate interval time series in heart health tracking.

## 6. Acknowledgments

This study was supported by Grant-in-aid for Young Scientists of the Japan Society for the Promotion of Science (JSPS), 2017-2019 and by Next Generation Interdisciplinary Research Project of NAIST.

